# Positive selection in multiple salivary gland proteins of Anophelinae reveals potential targets for vector control

**DOI:** 10.1101/2021.02.16.431392

**Authors:** Lucas Freitas, Mariana F. Nery

## Abstract

*Anopheles* is a genus belonging to the Culicidae family, which has great medical importance due to its role as a vector of *Plasmodium*, the parasite responsible for malaria. From *Anopheles’* functional genomics, great focus has been given to the salivary gland proteins (SGPs) group. This class of proteins is essential to blood-feeding behavior as they have attributes such as vasodilators and anti-clotting properties. Recently, a comprehensive review on *Anopheles* SGPs was performed, however the authors did not deeply explore the adaptive molecular evolution of these genes. In this context, this work aimed to perform a more detailed analysis of the adaptive molecular evolution of SGPs in *Anopheles*, carrying out positive selection and gene families evolution analysis on 824 SGPs. Our results show that most SGPs have positively selected sites that can be used as targets in the development of new strategies for vector control. Notably, we were not able to find any evidence of an accelerated shift in the copy-number variation of SGPs compared with other proteins, as suggested in previous works.

**Significance Statement:** Salivary gland proteins (SGPs) are essential to blood-feeding behavior in *Anopheles* and they are the most studied class of proteins in blood-feeding insects. However a proper molecular evolution analysis on SGPs in *Anopheles* is missing. In our analyses we observed that most SGPs have positively selected sites and we were not able to find any evidence of an accelerated shift in the copy-number of SGPs compared with other proteins, as stated in the literature. Our results can open new venues in the development of new strategies for vector control.

## Introduction

*Anopheles* is a genus belonging to the Culicidae family, which has great medical importance due to its role as a vector of *Plasmodium*, the parasite responsible for malaria. The impact of malaria in human well-being is substantial, with 228 million cases and 405,000 deaths reported in 2018, mainly in poor tropical countries (WHO 2019). Due to its medical importance, *Anopheles* mosquitoes are heavily-studied organisms with researches scattered in many scientific fields: behavior (Takken and Knols 1999), genomics (Neafsey *et al*. 2015), molecular biology (Bahia *et al*. 2013), and others. From *Anopheles’* functional genomics, great focus has been given to the salivary gland proteins (SGPs) group. This class of proteins is essential to blood-feeding behavior as they have attributes such as vasodilators and anti-clotting properties (Arcà and Ribeiro 2018).

Recently, Arcà *et al*. (2017) published a comprehensive review on *Anopheles* SGPs using data available from the 16 *Anopheles* genome project (Neafsey *et al*. 2015). They characterized and discussed each of the main SGPs found on these genomes from a comparative genomics point of view. However, regarding the adaptive molecular evolution of these genes, Arcà *et al*. (2017) only estimated the overall *dN*/*dS* (ω) ratio for each SGP. Although the ω ratio alone can give us a good overview of the evolution of a gene (e.g. a ω > 1 indicates that natural selection is favoring the fixation of new mutations, which we usually call positive selection), new approaches can provide more resolution on the impact of natural selection in molecular sequences, estimating ω for different lineages, sites, and sites in specific lineages (Yang 2019).

In this context, this work aimed to perform a more detailed analysis of the adaptive molecular evolution of SGPs in *Anopheles*. First, our results showed younger divergence time estimates within *Anopheles* lineages compared with previous works. Next, we observed that most SGPs have positively selected sites that can be used as targets in the development of new strategies for vector control. Notably, we were not able to find any evidence of an accelerated shift in the copy-number of SGPs compared with other proteins.

## Results and discussion

### Divergence time estimation

Recently, several works estimated the age of the last common ancestor of *Anopheles* between 80~110 million years ago (Ma) (Neafsey *et al*. 2015; Moreno *et al*. 2010; Freitas *et al*. 2015; Hao *et al*. 2017; Martinez-Villegas *et al*. 2019). In this work the split between *Anopheles* and *Aedes* was estimated at 130 Ma (Early Cretaceous) and the divergence of the three main subgenera was inferred at 72 Ma for *Nyssorhynchus* (Late Cretaceous-Paleogene), 53 Ma for *Anopheles*, and 39 Ma for *Cellia*, (both splits within the Paleogene). Further, the age of the gambiae complex was dated at 2.6 Ma, at the beginning of the Quaternary (Figure 1).

**Figure 1.**
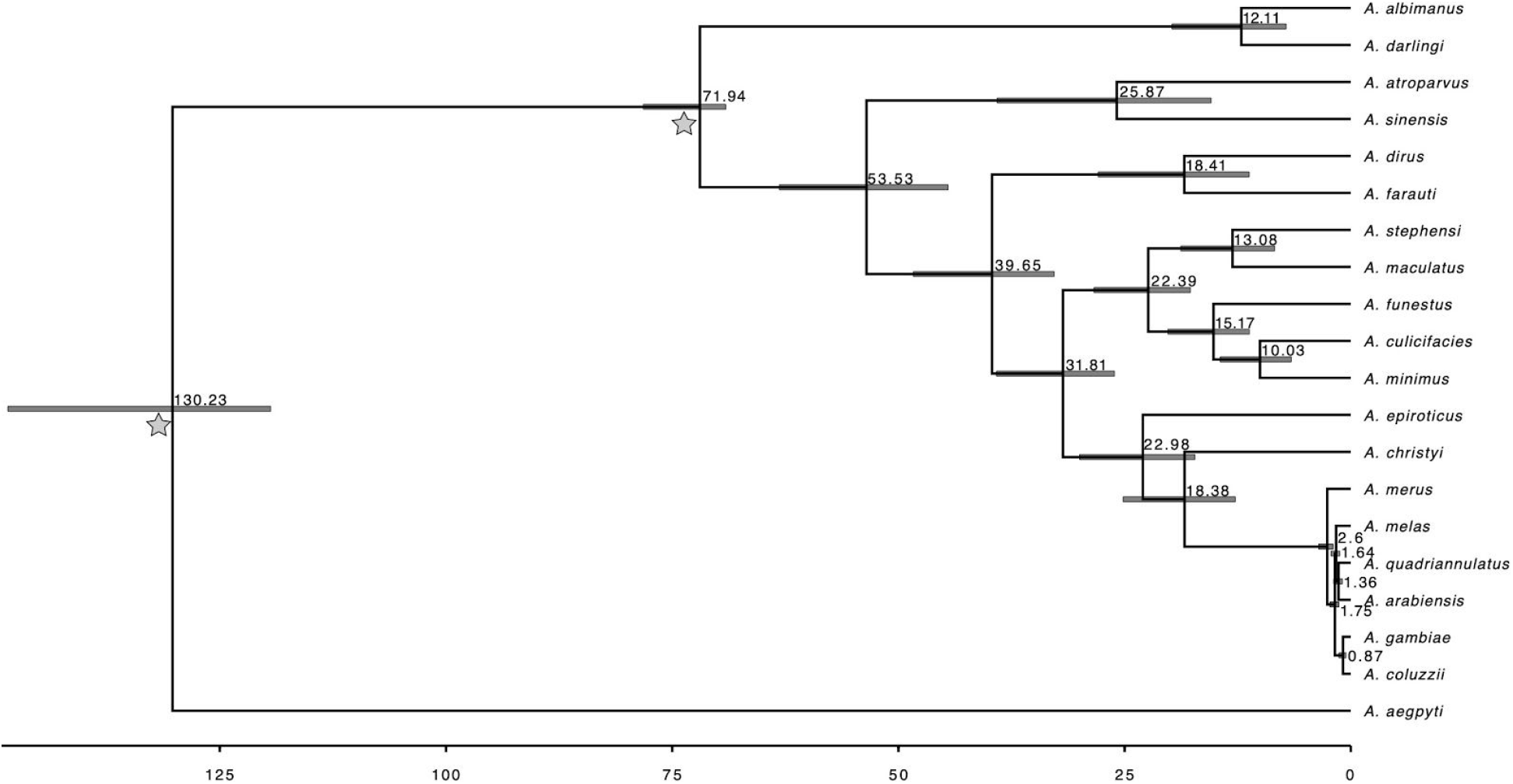
Phylogram of 19 *Anopheles* species used in this study. Numbers in the nodes represent the posterior mean age inferred by MCMCtree, while graybar scales are the 95% highest posterior density of the same analysis. The two gray stars represent the location of the calibration nodes used to estimate the divergence time and the x-axis is the timescale in million years ago.

Compared with Neafsey *et al*. (2015), the only other work that used a phylogenomic dataset, our divergence time estimates for these subgenera are younger: 53 Ma versus 80.3 Ma and 39 Ma versus 52.9 Ma for *Anopheles* and *Cellia*, respectively, however, many factors may influence the inference of divergence time as shown in other works (Warnock *et al*. 2015; Soares and Schrago 2015; Dos Reis, Zhu, and Yang 2014; Inoue, Donoghue, and Yang 2010).

### Adaptive molecular evolution of salivary gland proteins

In our analysis, OrthoFinder split the 824 SGPs into 37 gene families and after manual curation its final orthology had 821 sequences distributed in 33 gene families, nine more gene families when compared with Arcà *et al*. (2017). In OrthoFinder, D7 protein family was split into four gene families, hyp4.2/hyp13 was split into two gene families, hyp6.2/hyp8.2 was split into two gene families, and SG1 was split into five gene families.

To establish a comparison between both orthologies, all evolutionary analyses were performed twice, one for each orthology. We first screened the alignments to detect recombination breakpoints and these results were taken into account in the positive selection analysis.

Two distinct methodologies, MEME and FUBAR, were employed to identify episodic and pervasive positive selection, respectively. MEME was able to detect 119 sites evolving under episodic positive selection distributed in 26 out of 33 genes. Sixteen genes were shared between the two orthology assessment schemes, eight genes were exclusively found in OrthoFinder’s orthology, and only two genes were found only in orthologous from Arcà *et al*. (2017)’s orthology. (Table 1).

**Table 1.** Distribution of episodic and pervasive positively selected sites found in both orthologies schemes. Bold numbers are OrthoFinder exclusive orthologous; italic numbers are Arcà et al. (2017)’s exclusive orthologous; regular numbers are genes found in both orthology assessment schemes.

| Orthologous ID | Gene | Sites evolving under episodic positive selection | Sites evolving under pervasive positive selection |
| --- | --- | --- | --- |
| OG00 | Ag5 | 170, 220, 232, 322, 328, 336, 337, 407 | - |
| **OG01** | **D7r2** | **116, 185, 250, 253** | - |
| **OG02** | **SG1a** | **274, 316, 382, 419, 422, 580, 612, 624, 671** | - |
| OG03 | hyp37.7 | 502 | - |
| OG04 | Salivary Serine Protease | 93, 125, 126, 128, 205, 250, 265, 298, 302, 341, 377, 380 | - |
| OG05 | SG7 | 36, 80 | - |
| OG06 | Apyrase 5' nucleotidase | 196, 466 | 196 |
| OG07 | hyp10/hyp12 | 41, 62, 99 | - |
| OG08 | SG2 | 2, 7 | - |
| **OG09** | **D7L2** | **151, 205, 218, 222, 232, 252** | - |
| OG10 | hyp15/hyp17 | - | 26 |
| OG11 | 30kDa | - | 393/440 |
| **OG12** | **D7r5** | - | - |
| OG13 | Salivary peroxydase | - | - |
| OG14 | SG5 | 40, 43, 117, 145, 177, 354 | - |
| OG15 | SG8 | 323 | - |
| OG16 | SG9 | 226, 275, 433, 439, 460 | - |
| OG17 | cE5/anophelin | - | 28 |
| OG18 | Epoxy hydrolase | 13, 110, 133, 201, 305 | - |
| **OG19** | **hyp6.2** | - | - |
| **OG20** | **SG1-like2** | **177, 265, 452, 489** | - |
| **OG21** | **SG1b** | **240, 251, 262, 319, 353** | - |
| **OG22** | **TRIO (SG1)** | **185, 301, 366, 412, 474** | - |
| **OG23** | **D7L3** | **114, 198, 239** | **154** |
| OG24 | Salivary amylase | 43*, 63, 531/537, 825/827, 893/895, 969/971, 1061/1063, 1097/1099 | - |
| OG25 | Salivary maltase | 140, 307, 338, 463, 470, 571, 576, 647 | - |
| **OG26** | **SG1-like3** | **154, 248, 259, 375, 395** | - |
| OG27 | SG6 | 113 | - |
| OG28 | 55.3 kDa | 69, 73, 227, 282, 303, 427, 477, 497 | 73 |
| **OG29** | **hyp8.2** | - | - |
| OG30 | Salivary trypXII | 121/122, 124/125, 255/258, 337/340* | - |
| **OG31** | **Hyp4.2** | - | **19** |
| **OG32** | **Hyp13** | - | - |
| *Hyp4.2/Hyp13* | *Hyp4.2/Hyp13* | 19 | - |
| *SG1* | *SG1* | 962 | - |

\*Sites found only in Arcà *et al.* (2017)'s orthologous

FUBAR was able to detect seven sites evolving under pervasive positive selection with a posterior probability > 90% in seven distinct genes. Five of these seven genes were shared between both orthology assessment schemes and two of these genes were found only in OrthoFinder’s orthology. None gene was found in the four gene families exclusive of Arcà *et al*. (2017)’s orthology (Table 1). The discrepancy in the number of positively selected sites found between both methods is natural since MEME aims to detect episodic positive selection, while FUBAR aims to detect pervasive positive selection.

The joint analysis of MEME and FUBAR found just two positively selected sites in both methods: site 196 in Apyrase 5′ nucleotidase and site 73 in 55.3 kDa. Only 4/33 genes from OrthoFinder’s orthology did not show any site evolving under positive selection, either episodic or pervasive (OG000012/D7r5, OG000019/hyp6.2, OG000029/hyp8.2, and OG000032/hyp13).

Research on the molecular evolution of SGPs in Anophelinae is still scarce and aspects regarding patterns of molecular evolution are usually superficially discussed since they tend to be a small part of a bigger work of functional characterization of these SGPs. Further, several of these works lack a proper statistical analysis specifically designed to detect the presence of positive selection. For example, (Calvo *et al*. 2007, 2006; Valenzuela *et al*. 2003) imply that SGPs are evolving under positive selection because they are more diverse and/or their phylogeny have greater branch lengths compared with other proteins, but none formal positive selection test was performed.

Arcà *et al*. (2017) employed other evolutionary analysis and they estimated the ω ratio for SGPs showing that they are evolving at a faster rate compared with other proteins, however, they did not employ a statistical framework to detect individual sites subject to positive selection. Neafsey *et al*. (2015) estimated the ω ratio for SGPs obtaining the same results from Arcà *et al*. (2017), and they added other evolutionary analysis using site models to detect signatures of positive selection in individual sites. However, they did not report which sites were subjected to positive selection, an important result that can help us to understand the evolution of blood-feeding and also can lead to the development of new strategies for vector control (Anisimova 2015; Hammond *et al*. 2016).

The only work focused entirely on the molecular evolution of SGPs to date is Arcà *et al*. (2014). As the previous works mentioned here, they also found a high ω ratio for SGPs compared with other proteins, and they applied site models to detect signatures of positive selection in individual sites in SGPs, similarly to Neafsey *et al*. (2015). In their work, Arcà *et al*. (2014) were able to find eight sites in four genes evolving under pervasive selection and 22 sites in seven genes evolving under episodic selection. Although none of these sites were the same as those found in our work, this difference is normal as we used distinct softwares (or versions of the same software) and distinct datasets. The dataset from Arcà *et al*. (2014) is more focused on *A*. *gambiae* populational data, which is not suitable for methods designed for inter-species comparative genomics (Yang 2019) and contains fewer species than our work.

The inference of gene gains and losses for Anophelinae phylogeny using CAFE, estimated a λ rate of 0.0035 and 0.0032 for Arcà *et al*. (2017)’s orthology and OrthoFinder’s orthology, respectively. For both orthology schemes, none gene family shows evidence for rapid turnover rates and to our concern, no other work estimated a λ rate exclusively for SGPs in *Anophelinae*. Neafsey *et al*. (2015) estimated a λ rate of 0.0031 for all *Anophelinae* proteins, which is similar to the λ rate found in this work for SGPs.

Taken together, our results revealed that most SGPs (29 from 33) have many sites evolving under positive selection, similar to past works that have shown that SGPs have higher evolutionary rates compared with other proteins. The main differential of our work is that we were able to identify individual sites subjected to positive selection, which can open new avenues in new methodologies aiming to help vector control. Also, we showed that the λ rate of SGPs is similar to an overall λ rate for Anophelinae.

## Methodology

### Divergence time estimation and gene families evolution

We retrieved a total of 1,713 1:1 orthologous genes for 20 species of *Anopheles* and *Aedes aegypti* available at VectorBase (Giraldo-Calderón *et al*. 2015). We aligned each gene with PRANK v.170427 (Löytynoja and Goldman 2008) to infer individual gene trees using IQ-TREE v.2.0.4 (Minh *et al*. 2020).

Next, we used an Anophelinae phylogeny with species relationships based on Thawornwattana, Dalquen, and Yang (2018) for the *A*. *gambiae* complex and Neafsey *et al*. (2015) for the rest of the phylogenetic tree to select the 50 most clock-like genes using the SortaDate package (Smith, Brown, and Walker 2018). Then, the alignment of these 50 genes and the same Anophelinae phylogeny was used as input to MCMCtree v.4.9f (Yang 2007) to perform a Bayesian divergence time estimation with the approximated likelihood calculation described in (Dos Reis and Yang 2011). We used two calibration points based on the split of *Anopheles* and *Aedes* at 135 Ma (Freitas and Nery 2020) and the split between *A*. *gambiae* and *A*. *darlingi* at 79 Ma (Moreno *et al*. 2010).

### Adaptive molecular evolution of salivary gland proteins

We used 842 protein-coding sequences corresponding to 53 SGPs found in the *Anopheles* genus available at Arcà *et al*. (2017) and kindly provided by the authors. In Arcà *et al*. (2017) these 53 proteins were split into 24 gene families. To compare and to ensure the correct orthology, we used OrthoFinder v. 2.3.8 (Emms and Kelly 2019) to run a phylogenetic orthology assessment of these 842 protein-coding sequences. Therefore, all evolutionary tests were performed both in Arcà *et al*. (2017) orthology and in OrthoFinder’s orthology.

We used HyPhy package v. 2.5.14 (Pond *et al*. 2020) to perform three evolutionary analyses for each gene using GARD (Pond *et al*. 2006), MEME (Murrell *et al*. 2012), and FUBAR (Murrell *et al*. 2013). GARD screens a sequence alignment to identify recombination breakpoints and then it splits the sequence alignment into different partitions according to the recombination breakpoints. After screening for breakpoints we performed two positive selection tests.

Briefly, MEME and FUBAR aim to infer if the rate of non-synonymous substitutions (β) is greater than the rate of synonymous substitutions (α) at a specific codon site in the alignment. When β > α in a specific site, we assume this site is evolving under positive selection, i.e., natural selection is favoring new mutations that change the amino acid. The key difference between both methods is that MEME uses a mixed-effects maximum likelihood approach to detect *episodic* positive selection, i.e., to identify positively selected sites affecting a subset of lineages in the phylogeny, while FUBAR uses a Bayesian approach to detect *pervasive* positive selection, i.e., to identify positively selected sites across the whole phylogeny. We assumed a site to be positively selected if the *p*-value of MEME < 0.05 and the posterior probability of FUBAR > 90%.

To estimate the birth-death rate for SGPs we used CAFE v.5 (10.5281/zenodo.3625141). CAFE uses a stochastic birth and death model that takes into account phylogenetic branches lengths, duplication and deletion rates to infer the birth and death rate parameter (λ), which represents the average rate of genomic turnover (gains and losses) per gene per million years. As input for CAFE we used the timetree inferred above and a matrix with the number of genes per family per species.

## Acknowledgments

This work was supported by the Fundação de Amparo À Pesquisa do Estado de São Paulo [Grant numbers 2017/25058-2 and 2015/18269-1].

## Data Availability Statement

The sequences used in this study are the same from Arcà *et al*. (2017) and they were kindly provided by the authors. All scripts used in this work are available at https://github.com/freitas-lucas/SGP_Anopheles.

## References

Anisimova, Maria. 2015. “Darwin and Fisher Meet at Biotech: On the Potential of Computational Molecular Evolution in Industry.” BMC Evolutionary Biology 15 (May): 76.

Anisimova, Maria, Rasmus Nielsen, and Ziheng Yang. 2003. “Effect of Recombination on the Accuracy of the Likelihood Method for Detecting Positive Selection at Amino Acid Sites.” Genetics 164 (3): 1229–36.

Arcà, Bruno, Fabrizio Lombardo, Claudio J. Struchiner, and José M. C. Ribeiro. 2017. “Anopheline Salivary Protein Genes and Gene Families: An Evolutionary Overview after the Whole Genome Sequence of Sixteen Anopheles Species.” BMC Genomics 18 (1): 153.

Arcà, Bruno, and Josè M. C. Ribeiro. 2018. “Saliva of Hematophagous Insects: A Multifaceted Toolkit.” Current Opinion in Insect Science 29 (October): 102–9.

Arcà, B., C. J. Struchiner, V. M. Pham, G. Sferra, F. Lombardo, M. Pombi, and J. M. C. Ribeiro. 2014. “Positive Selection Drives Accelerated Evolution of Mosquito Salivary Genes Associated with Blood-Feeding.” Insect Molecular Biology 23 (1): 122–31.

Bahia, Ana C., José Henrique M. Oliveira, Marina S. Kubota, Helena R. C. Araújo, José B. P. Lima, Claudia Maria Ríos-Velásquez, Marcus Vinícius G. Lacerda, Pedro L. Oliveira, Yara M. Traub-Csekö, and Paulo F. P. Pimenta. 2013. “The Role of Reactive Oxygen Species in *Anopheles aquasalis* Response to *Plasmodium vivax* Infection.” PloS One 8 (2): e57014.

Calvo, Eric, Adama Dao, Van M. Pham, and José M. C. Ribeiro. 2007. “An Insight into the Sialome of *Anopheles funestus* Reveals an Emerging Pattern in Anopheline Salivary Protein Families.” Insect Biochemistry and Molecular Biology 37 (2): 164–75.

Calvo, Eric, Ben J. Mans, John F. Andersen, and José M. C. Ribeiro. 2006. “Function and Evolution of a Mosquito Salivary Protein Family.” The Journal of Biological Chemistry 281 (4): 1935–42.

Dos Reis, M., T. Zhu, and Z. Yang. 2014. “The Impact of the Rate Prior on Bayesian Estimation of Divergence Times with Multiple Loci.” Systematic Biology 63 (4): 555–65.

Dos Reis, M., and Z. Yang. 2011. “Approximate Likelihood Calculation on a Phylogeny for Bayesian Estimation of Divergence Times.” Molecular Biology and Evolution. https://doi.org/10.1093/molbev/msr045.

Emms, David M., and Steven Kelly. 2019. “OrthoFinder: Phylogenetic Orthology Inference for Comparative Genomics.” Genome Biology 20 (1): 238.

Freitas, Lucas A., Claudia A. M. Russo, Carolina M. Voloch, Olívio C. F. Mutaquiha, Lucas P. Marques, and Carlos G. Schrago. 2015. “Diversification of the Genus *Anopheles* and a Neotropical Clade from the Late Cretaceous.” PloS One 10 (8): e0134462.

Freitas, Lucas, and Mariana F. Nery. 2020. “Expansions and Contractions in Gene Families of Independently-Evolved Blood-Feeding Insects.” BMC Evolutionary Biology 20 (1): 1–8.

Giraldo-Calderón, Gloria I., Scott J. Emrich, Robert M. MacCallum, Gareth Maslen, Emmanuel Dialynas, Pantelis Topalis, Nicholas Ho, et al. 2015. “VectorBase: An Updated Bioinformatics Resource for Invertebrate Vectors and Other Organisms Related with Human Diseases.” Nucleic Acids Research 43 (Database issue): D707–13.

Hammond, Andrew, Roberto Galizi, Kyros Kyrou, Alekos Simoni, Carla Siniscalchi, Dimitris Katsanos, Matthew Gribble, et al. 2016. “A CRISPR-Cas9 Gene Drive System Targeting Female Reproduction in the Malaria Mosquito Vector *Anopheles gambiae*.” Nature Biotechnology 34 (1): 78–83.

Hao, You-Jin, Yi-Lin Zou, Yi-Ran Ding, Wen-Yue Xu, Zhen-Tian Yan, Xu-Dong Li, Wen-Bo Fu, Ting-Jing Li, and Bin Chen. 2017. “Complete Mitochondrial Genomes of *Anopheles stephensi* and *An. dirus* and Comparative Evolutionary Mitochondriomics of 50 Mosquitoes.” Scientific Reports. https://doi.org/10.1038/s41598-017-07977-0.

Inoue, Jun, Philip C. J. Donoghue, and Ziheng Yang. 2010. “The Impact of the Representation of Fossil Calibrations on Bayesian Estimation of Species Divergence Times.” Systematic Biology 59 (1): 74–89.

Löytynoja, Ari, and Nick Goldman. 2008. “Phylogeny-Aware Gap Placement Prevents Errors in Sequence Alignment and Evolutionary Analysis.” Science 320 (5883): 1632–35.

Martinez-Villegas, Luis, Juliana Assis-Geraldo, Leonardo B. Koerich, Travis C. Collier, Yoosook Lee, Bradley J. Main, Nilton B. Rodrigues, et al. 2019. “Characterization of the Complete Mitogenome of *Anopheles aquasalis*, and Phylogenetic Divergences among *Anopheles* from Diverse Geographic Zones.” PloS One 14 (9): e0219523.

Minh, Bui Quang, Heiko A. Schmidt, Olga Chernomor, Dominik Schrempf, Michael D. Woodhams, Arndt von Haeseler, and Robert Lanfear. 2020. “IQ-TREE 2: New Models and Efficient Methods for Phylogenetic Inference in the Genomic Era.” Molecular Biology and Evolution 37 (5): 1530–34.

Moreno, Marta, Osvaldo Marinotti, Jaroslaw Krzywinski, Wanderli P. Tadei, Anthony A. James, Nicole L. Achee, and Jan E. Conn. 2010. “Complete mtDNA Genomes of *Anopheles darlingi* and an Approach to Anopheline Divergence Time.” Malaria Journal. https://doi.org/10.1186/1475-2875-9-127.

Murrell, Ben, Sasha Moola, Amandla Mabona, Thomas Weighill, Daniel Sheward, Sergei L.Kosakovsky Pond, and Konrad Scheffler. 2013. “FUBAR: A Fast, Unconstrained Bayesian Approximation for Inferring Selection.” Molecular Biology and Evolution 30 (5): 1196–1205.

Murrell, Ben, Joel O. Wertheim, Sasha Moola, Thomas Weighill, Konrad Scheffler, and Sergei L. Kosakovsky Pond. 2012. “Detecting Individual Sites Subject to Episodic Diversifying Selection.” PLoS Genetics 8 (7): e1002764.

Neafsey, Daniel E., Robert M. Waterhouse, Mohammad R. Abai, Sergey S. Aganezov, Max A. Alekseyev, James E. Allen, James Amon, et al. 2015. “Mosquito Genomics. Highly Evolvable Malaria Vectors: The Genomes of 16 *Anopheles* Mosquitoes.” Science 347 (6217): 1258522.

Pond, Sergei L. Kosakovsky, Sergei L. Kosakovsky Pond, Art F. Y. Poon, Ryan Velazquez, Steven Weaver, N. Lance Hepler, Ben Murrell, et al. 2020. “HyPhy 2.5—A Customizable Platform for Evolutionary Hypothesis Testing Using Phylogenies.” Molecular Biology and Evolution. https://doi.org/10.1093/molbev/msz197.

Pond, S. L. Kosakovsky, S. L. Kosakovsky Pond, D. Posada, M. B. Gravenor, C. H. Woelk, and S. D. W. Frost. 2006. “GARD: A Genetic Algorithm for Recombination Detection.” Bioinformatics. https://doi.org/10.1093/bioinformatics/btl474.

Smith, Stephen A., Joseph W. Brown, and Joseph F. Walker. 2018. “So Many Genes, so Little Time: A Practical Approach to Divergence-Time Estimation in the Genomic Era.” PloS One 13 (5): e0197433.

Soares, André E. R., and Carlos G. Schrago. 2015. “The Influence of Taxon Sampling on Bayesian Divergence Time Inference under Scenarios of Rate Heterogeneity among Lineages.” Journal of Theoretical Biology 364 (January): 31–39.

Takken, W., and B. G. Knols. 1999. “Odor-Mediated Behavior of Afrotropical Malaria Mosquitoes.” Annual Review of Entomology 44: 131–57.

Thawornwattana, Yuttapong, Daniel Dalquen, and Ziheng Yang. 2018. “Coalescent Analysis of Phylogenomic Data Confidently Resolves the Species Relationships in the *Anopheles gambiae* Species Complex.” Molecular Biology and Evolution 35 (10): 2512–27.

Valenzuela, Jesus G., Ivo M. B. Francischetti, Van My Pham, Mark K. Garfield, and José M. C. Ribeiro. 2003. “Exploring the Salivary Gland Transcriptome and Proteome of the *Anopheles stephensi* Mosquito.” Insect Biochemistry and Molecular Biology 33 (7): 717–32.

Warnock, Rachel C. M., James F. Parham, Walter G. Joyce, Tyler R. Lyson, and Philip C. J. Donoghue. 2015. “Calibration Uncertainty in Molecular Dating Analyses: There Is No Substitute for the Prior Evaluation of Time Priors.” Proceedings. Biological Sciences / The Royal Society 282 (1798): 20141013.

World Health Organization. 2019. World Malaria Report 2019.

Yang, Ziheng. 2007. “PAML 4: Phylogenetic Analysis by Maximum Likelihood.” Molecular Biology and Evolution 24 (8): 1586–91.

Yang, Ziheng. 2019. “Adaptive Molecular Evolution.” In Handbook of Statistical Genomics, edited by David Balding Ida Moltke, 369–96. Wiley.

